# Does dopamine synthesis capacity predict individual variation in curiosity?

**DOI:** 10.1101/2020.10.13.337477

**Authors:** Lieke L. F. van Lieshout, Ruben van den Bosch, Lieke Hofmans, Floris P. de Lange, Roshan Cools

## Abstract

Curiosity, which can be defined as “intrinsically motivated information-seeking”, is an important driving force in our everyday lives. Based on previous evidence demonstrating a link between information prediction errors and dopamine neuronal firing rates, we asked whether the drive to seek information varies with individual differences in dopamine synthesis capacity. In order to investigate this, we let participants perform a lottery task in which we independently manipulated outcome uncertainty, outcome valence (gains versus losses) and expected value, and asked participants to indicate their curiosity for each presented lottery. In a separate session, participants underwent an [^18^F]DOPA PET scan to quantify their dopamine synthesis capacity. We replicate previous behavioral results, showing that curiosity is a function of outcome uncertainty as well as outcome valence (gain versus loss). However, we found no evidence that curiosity or the sensitivity to outcome uncertainty, outcome valence and expected value was related to participants’ dopamine synthesis capacity in the ventral striatum, the caudate nucleus or the putamen. These findings stress the need for further studies into the role of dopamine in (different types of) curiosity.

## Introduction

Curiosity is pervasive in our everyday lives and humans devote a substantial part of their time seeking and consuming information. On occasion, this information is directly relevant to us. We probably all remember the times when we were studying the information in our text books to increase the probability of passing our exam. In this type of situation, processing the information we encounter is directly relevant for achieving our goals or to obtain higher rewards (i.e. a higher grade for your exam). We call this type of information seeking “instrumental curiosity” or “goal-directed information seeking” (e.g. Addicott, Pearson, Sweitzer, Barack, & Platt, 2017; Averbeck, 2015; Daw & Doya, 2006). However, curiosity appears to be a broad feature, which also generalizes to situations in which information cannot inform action or directly increase our rewards (Bromberg-Martin & Hikosaka, 2009, 2011; Charpentier, Bromberg-Martin, & Sharot, 2018; Kobayashi, Ravaioli, Baranès, Woodford, & Gottlieb, 2019; van Lieshout, Vandenbroucke, Müller, Cools, & de Lange, 2018). When we are curious about information that serves no direct purpose, we refer to this as “non-instrumental curiosity” (see also Kidd & Hayden, 2015).

In fact, humans and other animals show a strong drive for information (Kreps & Porteus, 1978; Lieberman, Cathro, Nichol, & Watson, 1997; Prokasy, 1956). For example, studies with macaque monkeys have revealed a preference for receiving information about upcoming primary rewards, even though the information did not alter the likelihood of actually receiving the reward (Bromberg-Martin & Hikosaka, 2009, 2011). They were even willing to give up a substantial proportion of their reward in order to receive this advance information (Blanchard, Hayden, & Bromberg-Martin, 2015). Information and primary reward have been demonstrated to implicate the same neural structures: midbrain dopamine (DA) neurons and lateral habenula (LHb) neurons coding for reward prediction errors (i.e. the difference between expected and received reward) also coded for information prediction errors, i.e. the difference between expected information and received information (Bromberg-Martin & Hikosaka, 2009, 2011). In addition to this preference for information, prior work has demonstrated that humans show a preference for positive versus negative belief updating (Charpentier et al., 2018; van Lieshout, Traast, de Lange, & Cools, 2019). In a recent study, Charpentier and colleagues (2018) demonstrated that neural activity in the orbitofrontal cortex (OFC) signals the desire to gain knowledge over ignorance (see also Blanchard et al., 2015; van Lieshout et al., 2018), regardless of valence. However, activity in the mesolimbic reward circuitry (VTN/SN) was modulated by the opportunity to gain knowledge about positive, but not negative outcomes. As such, it might be the case that dopamine modulates the desire to seek information, even in non-instrumental contexts. However, no direct evidence for this link exists in humans.

Inspired by the well-established link between dopamine neuron firing and reward prediction errors (Schultz, Dayan, & Montague, 1997), previous work linked dopamine synthesis capacity to ventral striatal coding of reward prediction errors (Deserno et al., 2015; Schlagenhauf et al., 2013; Boehme et al., 2015), reward-based reversal learning (Cools et al., 2009), cognitive control (Aarts et al., 2014) and cognitive effort (Hofmans, Papadopetraki, et al., 2020; Westbrook, van den Bosch, et al., 2020). Here we asked whether individual variability in a different form of cognitive motivation, namely non-instrumental curiosity, is also linked to variation in dopamine synthesis capacity. This question is based on prior work by Bromberg-Martin and Hikosaka (2009, 2011) and Charpentier and colleagues (2018), who demonstrated that the desire for knowledge in non-instrumental settings implicates the mesolimbic reward circuitry. We investigated to what extent curiosity and the motives underlying human curiosity, such as outcome uncertainty (Kobayashi et al., 2019; Romero Verdugo, van Lieshout, de Lange, & Cools, 2020; van Lieshout, de Lange, & Cools, 2020a; van Lieshout et al., 2018; van Lieshout, Traast, et al., 2019), outcome valence (Charpentier et al., 2018; Marvin & Shohamy, 2016; van Lieshout et al., 2020a, van Lieshout, Traast, et al., 2019) and expected value (Charpentier et al., 2018; Kobayashi et al., 2019; Romero Verdugo et al., 2020; van Lieshout, Traast, et al., 2019), correlate with an individual’s dopamine synthesis capacity, measured with [^18^F]DOPA positron emission tomography (PET).

### The present experiment

We adapted a lottery task we used previously (van Lieshout et al., 2020a, 2018; van Lieshout, Traast, et al., 2019). In this task, every trial is a lottery consisting of a vase containing a mix of red and blue marbles, which were associated with monetary values. One marble would be randomly selected from every vase and participants would gain or lose the money associated with the marble. Participants had to indicate their curiosity about the outcome of a presented lottery, while they were clearly instructed that the information provided by the outcome was non-instrumental: all outcomes were obtained regardless of participants’ curiosity decisions and they had no way of influencing how much they would gain or lose during the task. This task enabled the independent manipulation of the uncertainty of the lottery outcomes, the outcome valence (whether the lottery contained gains or losses), and the amount of these gains and losses (expected value).

In a separate session, participants underwent an [^18^F]DOPA PET scan to quantify their dopamine synthesis capacity. We investigated whether participants’ dopamine synthesis capacity predicts the extent to which participants are curious about the outcomes, perhaps as a function of outcome uncertainty, outcome valence (gain/loss) and, potentially, expected value (Charpentier et al., 2018; Kobayashi et al., 2019; Romero Verdugo et al., 2020; van Lieshout et al., 2020a; van Lieshout, de Lange, & Cools, 2020b; van Lieshout et al., 2018; van Lieshout, de Lange, & Cools, 2019; van Lieshout, Traast, et al., 2019). For each of these factors, the null hypothesis was that the extent to which people are curious, and show sensitivity to the effects that drive curiosity, would be independent of their dopamine synthesis capacity. The alternative hypotheses were that participants with higher dopamine synthesis capacity would be [1] overall more curious, [2] more curious about gains versus losses, [3] more curious about high versus low outcome uncertainty and/or [4] more curious about high versus low expected value than participants with lower dopamine synthesis capacity.

## Methods

### Preregistration and data & code availability

The experiment and its analyses were preregistered on the Open Science Framework (osf.io/9svtg/). All data and code used for stimulus presentation and analyses will be available on the Donders Repository.

### Participants

Forty-five out of a total of 94 participants in a previous [^18^F]DOPA PET study (protocol NL57538.091.16; trial register NTR6140, www.trialregister.nl/trial/5959, see also Hofmans, Papadopetraki, et al., 2020; Westbrook, van den Bosch, et al., 2020) accepted the invitation to participate in the current experiment. The time between the PET scan and the current experiment ranged between 0.3 and 1.8 years (mean = 1.0; SD = 0.4). In the session containing the current experiment, the participants performed [1] a Stroop task (see Hofmans et al., 2020), [2] an oddball detection task, [3] the current experiment and [4] were asked to install a smartphone application tracking their screen touches (see Westbrook, Ghosh, van den Bosch, & Cools, 2020).

All participants of the current experiment were Dutch native speakers and right-handed. Following the preregistration, one participant was excluded because of too many missed trials (missed > 10% of all trials). Another participant was excluded due to a lack of variation in responses (gave the same curiosity rating on every trial). Although the lack of variation in responses was not a preregistered criterion, we decided to exclude the participant because no models could be fitted on the participant’s data and we could not be certain that the participant was engaged with the task. As a result, the final sample of the current experiment consisted of forty-three participants (22 women, age 23.5 ± 4.78, mean ± SD). The participants gave written informed consent according to the declaration of Helsinki prior to participation. The experiment was approved by the local ethics committee (CMO Arnhem-Nijmegen, The Netherlands) under a general ethics approval protocol (“Imaging Human Cognition”, CMO 2014/288) and was conducted in compliance with these guidelines.

### Behavioral paradigm

The experiment consisted of a lottery task (Figure 1). Each trial started with an image of a vase containing twenty marbles, each of which could be either red or blue. The vases could be configured in two possible ways: (1) 75%-25% vases: 15 marbles had one color and 5 marbles the other color, (2) 50%-50% vases: 10 marbles had one color and 10 marbles the other color. Both colored marbles were associated with a monetary value that participants could either gain or lose. These monetary values varied on a trial-by-trial basis between +10 and +90 cents in gain trials and between −90 and −10 cents in loss trials (both in steps of 10 cents). All combinations of monetary values associated with red and blue marbles were possible, except for combinations of the same monetary values.

**Figure 1.**
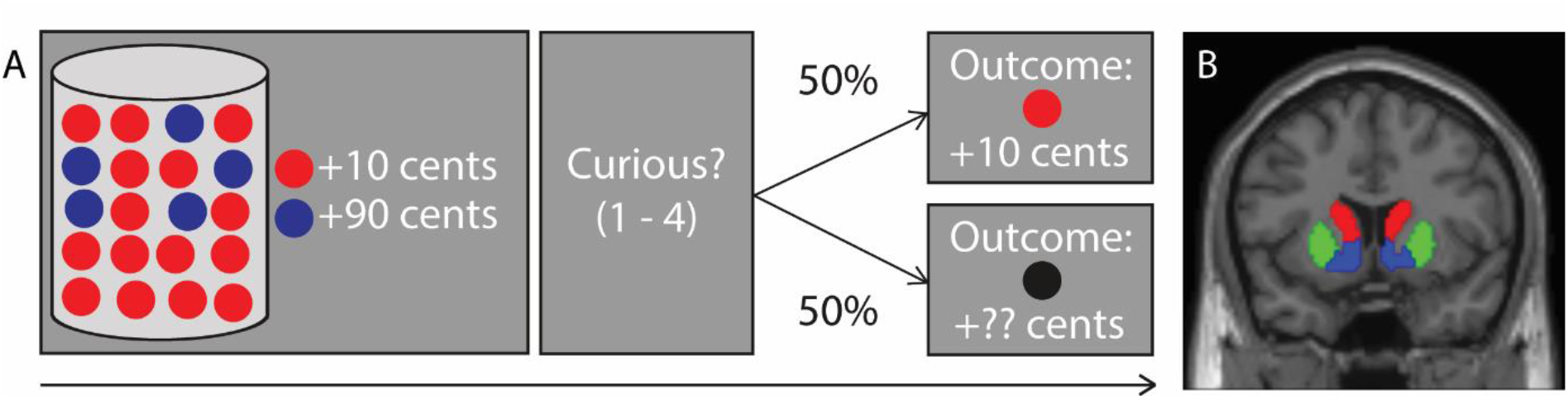
Schematic depiction of the task and the regions of interest. **A.** Schematic depiction of a gain trial in the lottery task. Participants saw a screen on which a vase with a mix of 20 red and blue marbles was presented and the monetary values associated with the colored marbles. These values could either both be positive (in gain trials as depicted here) or both be negative (in the loss trials). Participants were told that one of the marbles would be randomly selected from the vase and that they would be awarded the money associated with this marble. Next, participants indicated how curious they were about seeing the outcome (1 – 4). There was a 50% chance of seeing the outcome, regardless of the participants’ curiosity response. Importantly, a marble was randomly selected on every trial and participants were instructed that they were awarded the money associated with this marble, also if they would not see the outcome of a trial. See Methods (*Behavioral paradigm*) for details on the timing of the experiment. **B.** Coronal view of the regions of interest: the ventral striatum (blue), the caudate nucleus (red) and the putamen (green).

The participants were informed that on each trial, one marble would be selected from the vase and that they would gain or lose the money associated with the selected marble. First, the participants saw the vase, the marbles and the monetary values associated with the marbles (3s), followed by a blank screen (0.5s) and a response screen. On the response screen, participants could indicate how curious they were about the lottery outcome (“How curious are you about the outcome?”) on a scale from 1 – 4 using a button box. They had to use their right index finger, middle finger, ring finger and little finger to give curiosity ratings of 1, 2, 3 and 4 respectively. This response screen was presented until the participant gave a response, with a limit of 2.5s. The response screen was followed by a blank screen (0.5s), and an outcome screen (2s). When the outcome was presented, the outcome screen depicted the vase, the marbles and monetary values associated with the marbles again, together with a box in which they saw the colored marble that was selected and the amount of money they gained or lost in that trial. When the outcome was not presented, the participants saw a black marble instead of a colored marble and question marks at the location of the monetary value. After a trial ended, there was a blank screen (1s), after which the next trial started.

Participants were informed that they had a 50% chance of seeing the outcome of a particular trial and that they could not influence whether the outcome would be presented or not. This manipulation was explicitly instructed to subjects and it uncoupled curiosity responses from the actual receipt of the outcome. Additionally, they were explicitly instructed that they could not influence which marble would be selected and how much money they would gain or lose. However, they were made aware that a marble would be randomly selected on every trial and that they would actually gain or lose the money associated with that marble, regardless of outcome presentation. The money they won or lost in every trial would be summed and the sum of money would be added to or subtracted from the money they earned for participation. In the end, the task was set up in a way that participants would always receive a bonus of 50 cents on top their base payment for participation.

For both experiments, the participants completed a total of 144 trials (72 gain trials and 72 loss trials). In turn, each vase configuration was presented on 72 occasions (36 times for gain trials and 36 times for loss trials). The trials were divided in 2 blocks of 72 trials. The trials were pseudo-randomized, such that participants were never presented with more than 4 gain trials or 4 loss trials in a row. The experiment lasted ~ 25 minutes in total.

### PET acquisition

In a separate session, all participants underwent an [^18^F]DOPA PET scan to quantify dopamine synthesis capacity. Uptake of the radiotracer [^18^F]DOPA indexes the rate at which dopamine is synthesized in (the terminals of) midbrain dopamine neurons, providing a relatively stable trait index of dopamine function (Egerton, Demjaha, McGuire, Mehta, & Howes, 2010). The PET scans were acquired on a PET/CT scanner (Siemens Biograph mCT; Siemens Medical Systems, Erlangen, Germany) at the Department of Nuclear Medicine of the Radboudumc, using an [^18^F]DOPA radiotracer, produced by the Radboud Translational Medicine department. Participants received 150mg of carbidopa and 400mg of entacapone 50 minutes before scanning to minimize peripheral metabolism of [^18^F]DOPA by decarboxylase and COMT, respectively, thereby increasing signal to noise ratio in the brain. After a bolus injection of [^18^F]DOPA (185MBq; approximately 5mCi) into the antecubital vein, a dynamic PET scan was collected over 89 minutes and divided into 24 frames (4×1, 3×2, 3×3, 14×5 min).

### Structural MRI

A high-resolution anatomical scan, T1-weighted MP-RAGE sequence (repetition time = 2300 ms, echo time = 3.03 ms, 192 sagittal slices, field of view = 256 mm, voxel size 1 mm isometric) was acquired using a Siemens 3T MR scanner with a 64-channel coil. These were used for coregistration and spatial normalization of the PET scans.

### PET analysis

The PET data (4 × 4 × 3 mm voxel size; 5mm slice thickness; 200 × 200 × 75 matrix) were reconstructed with weighted attenuation correction and time-of-flight recovery, scatter corrected, and smoothed with a 3mm full-width-at-half-maximum (FWHM) kernel. After reconstruction, the PET data were preprocessed and analysed using SPM12 (http://www.fil.ion.ucl.ac.uk/spm/). All PET frames were realigned to the mean image, and then coregistered to the anatomical MRI scan, using the mean PET image of the first 11 frames, which has a better range in image contrast outside the striatum than a mean image over the whole scan time. Dopamine synthesis capacity was computed per voxel as [^18^F]DOPA influx constant per minute (Ki) relative to the cerebellar grey matter reference region, using Gjedde-Patlak graphical analysis on the PET frames from the 24th to 89th minute (Patlak, Blasberg, & Fenstermacher, 1983). The individual cerebellum grey matter masks were obtained by segmenting the individuals’ anatomical MRI scan, using Freesurfer (https://surfer.nmr.mgh.harvard.edu/). The resulting individual Ki maps were spatially normalized and smoothed using an 8mm FWHM kernel (see also Hofmans, Papadopetraki, et al., 2020; Westbrook, van den Bosch, et al., 2020)

The Ki values were extracted from masks defining regions of interest based on an independent, functional connectivity-based parcellation of the striatum (Piray, den Ouden, van der Schaaf, Toni, & Cools, 2015). In particular, we extracted Ki values from 3 striatal regions (Figure 1B) – the ventral striatum / nucleus accumbens (607 voxels), the caudate nucleus (817 voxels) and the putamen (1495 voxels). We averaged across all voxels in each region for individual difference analyses. As such, we obtained 3 Ki values for each participant, one for each striatal region of interest.

### Experimental design

We investigated main effects of outcome valence (gain/loss), outcome uncertainty and (absolute) expected value on curiosity ratings. Additionally, we assessed whether the effects of outcome uncertainty and absolute expected value on curiosity differed between gain and loss trials. This was done by assessing the significance of the interaction effects between outcome uncertainty and outcome valence (gain/loss), and absolute expected value and outcome valence (gain/loss) on curiosity.

Next, we investigated whether there was a main effect of dopamine synthesis capacity on curiosity and whether the effects of outcome valence (gain/loss), outcome uncertainty and (absolute) expected value on curiosity varied with dopamine synthesis capacity. This was done by assessing the significance of the interaction effects between dopamine synthesis capacity (Ki values) and outcome valence (gain/loss), between dopamine synthesis capacity and outcome uncertainty, between dopamine synthesis capacity and absolute expected value on the curiosity ratings, and between dopamine synthesis capacity, absolute expected value and outcome valence (gain/loss) on the curiosity ratings.

In order to do so, a value of outcome uncertainty and expected value was calculated for every trial (X) as follows:

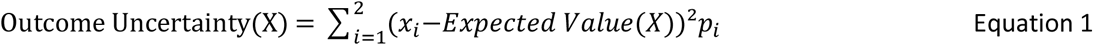

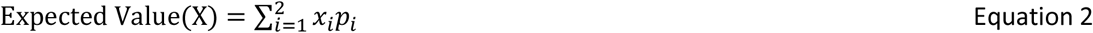

where x_i_ is the monetary value associated with marble (i), and pi the probability that this marble will be drawn. Hereby, outcome uncertainty (Equation 1) reflects the spread of the possible outcomes in trial (X), and expected value (Equation 2) reflects the mean expected reward contained in trial (X). It should be noted that we used a different calculation for outcome uncertainty in one of our previous studies (van Lieshout et al., 2018), and we initially preregistered to use that calculation of outcome uncertainty in the current manuscript as well. However, both metrics are almost identical, and variance is a more common measure of uncertainty (see for example Preuschoff, Quartz, & Bossaerts, 2008; Symmonds, Wright, Bach, & Dolan, 2011). Therefore we decided to operationalize outcome uncertainty as variance instead.

Expected value reflects the mean expected value of reward contained in a trial. Note that the expected value is always positive in a gain trial and always negative in a loss trial. To compare the effects of expected value between the gain and loss trials, we used absolute expected value in the analyses, such that − 90 cents in a loss trial is treated the same as + 90 cents in a gain trial etc. However, the metric of interest here is the effect of reward magnitude (signed expected value) on curiosity, which is reflected in the interaction between outcome valence (gain/loss) and absolute expected value.

### Primary statistical analyses

As preregistered, the data were analysed using a combination of mixed effects modelling in R (R Core Team, 2013; RRID:SCR_001905) and more classical repeated measures ANOVAs in SPSS (RRID:SCR_002865). The results of the mixed effects modelling in R are reported in the main text and the results of the repeated measures ANOVA are reported in Supplement 1. This allows the reader to verify the robustness of the results and demonstrate that our conclusions do not depend on the analytical framework employed (see also van Lieshout, Traast, et al., 2019; van Lieshout, de Lange, & Cools, 2020a).

We performed the analyses using the brm function of the BRMS package (Bürkner, 2017) in R. The analyses were performed as preregistered, except that we also added the three-way interaction between “dopamine synthesis capacity (Ki value)”, “absolute expected value” and “outcome valence (gain/loss)” to the models. We ran three main models for three striatal regions of interest separately (i.e. the ventral striatum, the caudate nucleus and the putamen). The main models included “curiosity rating” as an ordinal dependent variable. The main effects of “dopamine synthesis capacity (Ki value)”, “outcome valence (gain/loss)”, “outcome uncertainty” and “absolute expected value” were included as fixed effects. Additionally, the main models included interaction effects between “dopamine synthesis capacity (Ki value)” and “outcome valence (gain/loss)”, between “dopamine synthesis capacity (Ki value)” and “outcome uncertainty”, between “dopamine synthesis capacity” and “absolute expected value”, between “dopamine synthesis capacity”, “absolute expected value” and “outcome valence (gain/loss)”, between “outcome valence (gain/loss)” and “outcome uncertainty” and between “outcome valence (gain/loss)” and “absolute expected value” as fixed effects. A random intercept and random slopes for the main effects of “outcome valence (gain/loss)”, “outcome uncertainty” and “absolute expected value”, as well as for the interaction effects between “outcome valence (gain/loss)” and “outcome uncertainty” and between “outcome valence (gain/loss)” and “absolute expected value” were included per participant.

The predictors for dopamine synthesis capacity (Ki value), outcome uncertainty and absolute expected value were mean centered and scaled. We used the default priors of the brms package (Cauchy priors and LKJ priors for correlation parameters; Bürkner, 2017). The main models were fit using four chains with 10000 iterations each (5000 warm up) and inspected for convergence. Coefficients were deemed statistically significant if the associated 95% posterior credible intervals did not contain zero.

If any of the interaction effects with “outcome valence (gain/loss)” was significant in the main model, we used the brm function of the BRMS package (Bürkner, 2017) to model the gain and loss trials separately. All other conventions were as described for the main models, except that all effects including “outcome valence (gain/loss)” were omitted from the models.

Additionally, we ran a similar model that was set up in the same way as described above, except that we did not include the main and interaction effects with dopamine synthesis capacity (Ki value). Initially, we did not preregister to run this additional model, but we did so to provide one independent estimation of the task effects, instead of three results per task effect originating from the three models described above. It should be noted that the results are essentially the same when adding either one of the three Ki values to the model.

## Results

### Main task effects

We found a main effect of outcome uncertainty, such that participants were more curious about high compared with low outcome uncertainty (**BRMS:** 95% CI [1.43, 2.06]). In addition, we found a main effect of outcome valence, such that participants were more curious about gains compared with losses (**BRMS:** 95% CI [.15, .74]). However, there was no interaction between outcome uncertainty and outcome valence (**BRMS:** 95% CI [−.06, .13]), indicating that these effects operate independently (Figure 2; van Lieshout, et al, 2020a; van Lieshout, Traast, et al., 2019).

**Figure 2.**
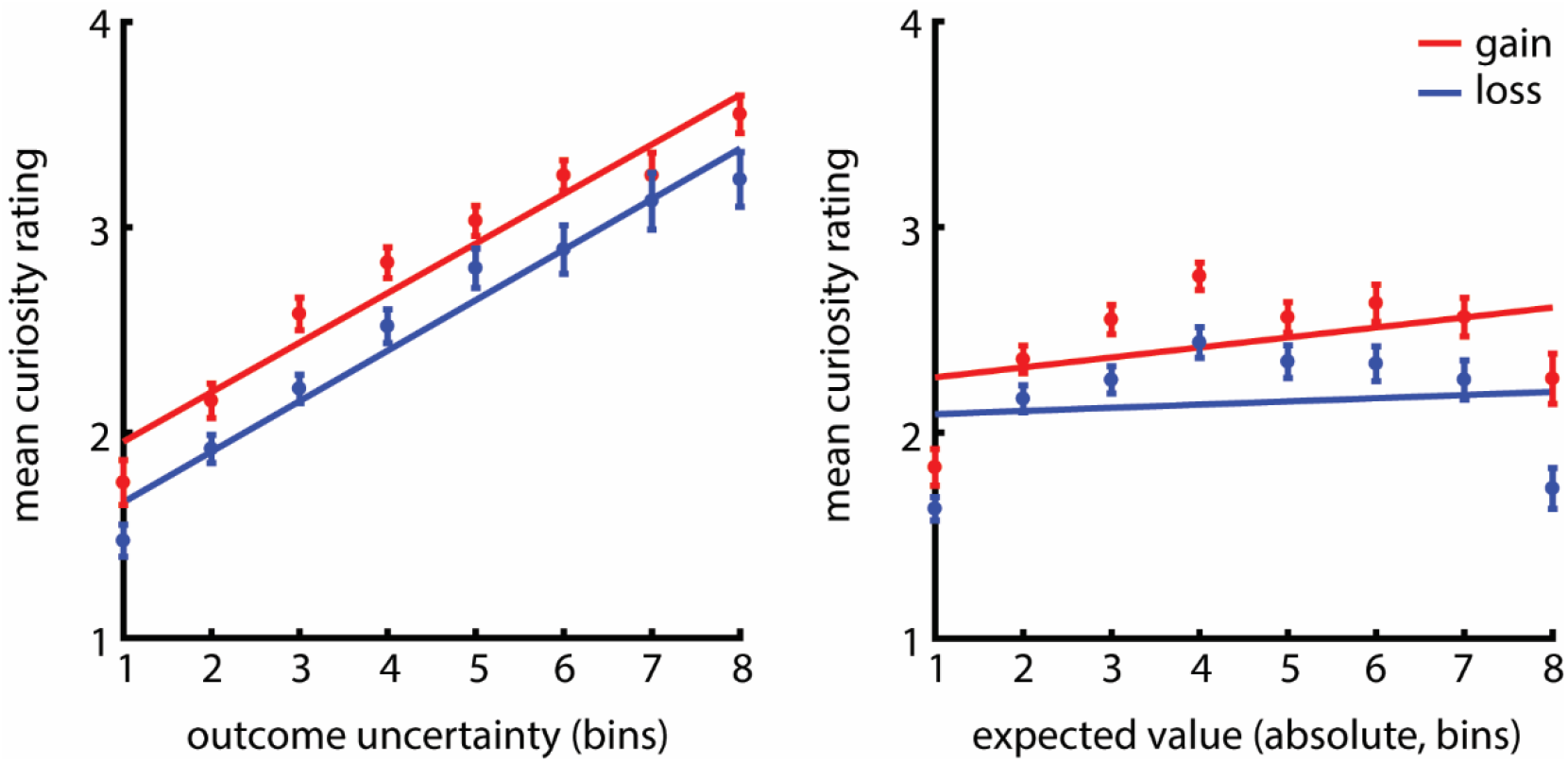
Behavioral results of the experiment. The x-axis depicts percentile bins of the values of outcome uncertainty (left) and the absolute expected values (right) for gains (in red) and losses (in blue). To this end, we divided the levels of outcome uncertainty in eight percentile bins per gain/loss condition, such that the 1st bin represents 1/8th of the lowest levels of outcome uncertainty, the 2nd bin represents the 1/8th – 2/8th of the lowest levels, etc. The absolute expected values were also divided in eight percentile bins, such that the 1st bin represents 1/8th of the lowest values, the 2nd bin the 1/8th – 2/8th of the lowest values, etc. The y-axis depicts the mean curiosity rating for each percentile bin of outcome uncertainty and absolute expected value for gains and losses. The error bars depict the standard error of the mean (SEM) and least square lines illustrate the effects. The experiment showed that curiosity was higher for gains than for losses and that there was a monotonic increase of curiosity with outcome uncertainty. In addition, curiosity increased with the amount of money that could be gained, but there was no effect of absolute expected value on curiosity in loss trials.

There was also a main effect of absolute expected value (**BRMS:** 95% CI [.07, .31]), and evidence for an interaction between outcome valence and absolute expected value (**BRMS:** 95% CI [.009, .20]). However, the interaction effect appears to be less robust and was not replicated when analyzing the data with repeated measures ANOVAs (Supplement and Supplementary Figure 1). When analyzing the gain and loss trials separately, we found that participants were more curious about higher compared with lower gains (**BRMS:** 95% CI [.15, 44]), but that there was no difference between whether participants would lose more or less money (**BRMS:** 95% CI [−.08, .25]).

### Individual differences

Next, we investigated whether the effects of outcome valence (gain/loss), outcome uncertainty, absolute expected value and the interaction between outcome valence and absolute expected value on curiosity varied with individual differences in dopamine synthesis capacity (Ki values) in the striatal regions of interest (ventral striatum, caudate nucleus and putamen; Figure 3 and Figure 4).

**Figure 3.**
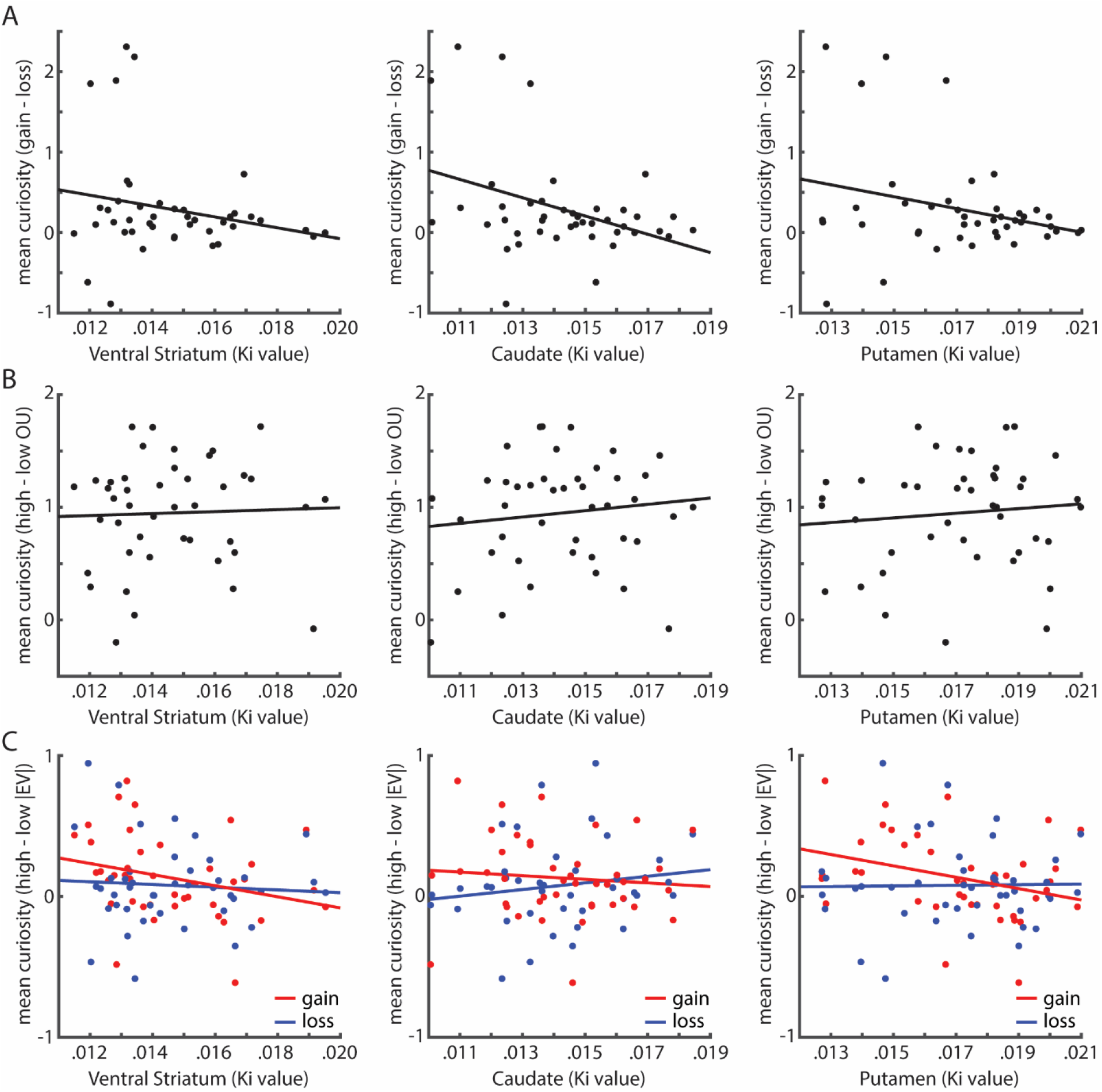
Individual differences. **A.** To visualize the extent to which sensitivity to gain versus loss trials depended on dopamine synthesis capacity, we calculated the difference between mean curiosity in gain trials and mean curiosity in loss trials per participant. These differences were plotted against the dopamine synthesis capacity (Ki values) for the three striatal regions of interest separately. **B.** The same was done for the effect of outcome uncertainty by calculating the difference between mean curiosity for high outcome uncertainty and low uncertainty per participant. **C.** The same was done for the effect of absolute expected value for gains and losses separately. We calculated the difference between mean curiosity for high absolute expected value and low absolute expected value for gains (in red) and losses (in blue) per participant.

**Figure 4.**
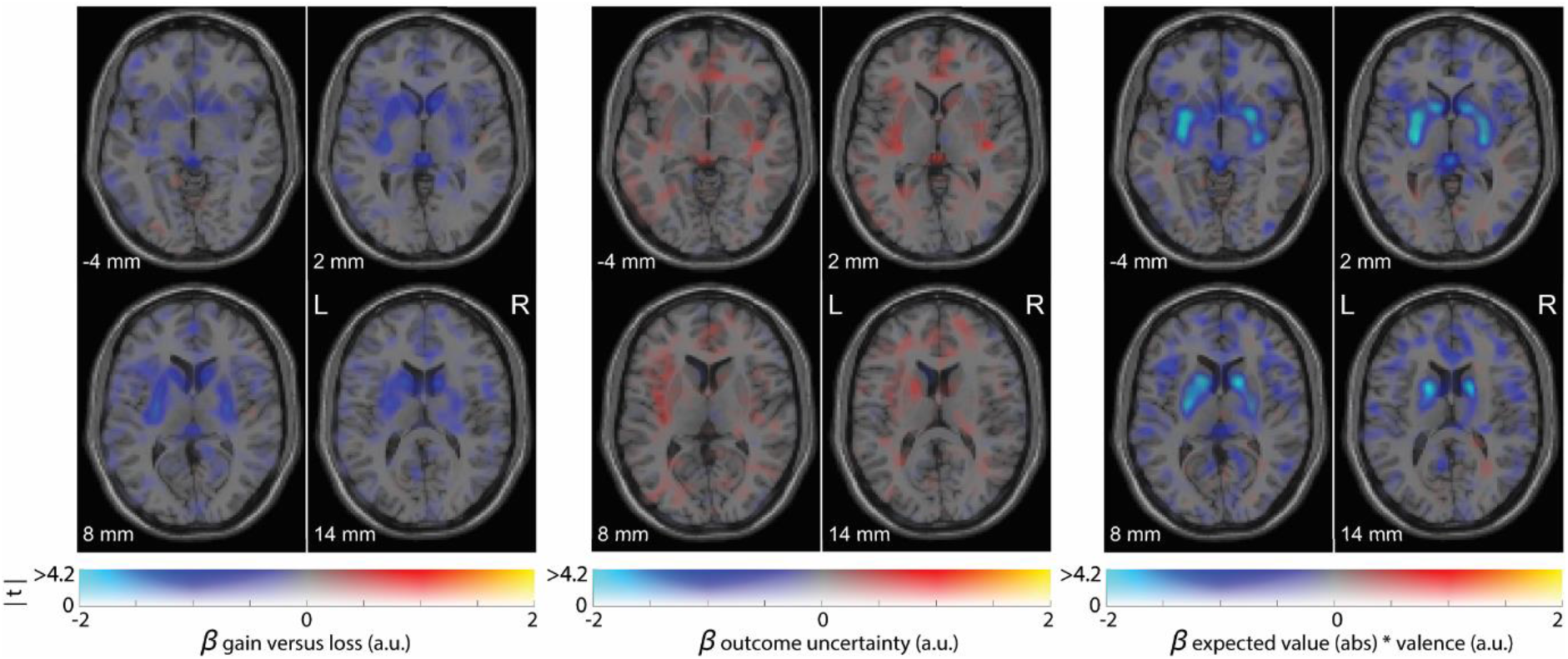
Association of dopamine synthesis capacity with the task effects. Figure 4 depicts the association of dopamine synthesis capacity with the outcome valence (gain/loss) effect, the outcome uncertainty effect and the interaction effect between absolute expected value and outcome valence (gain/loss). Voxels show a positive (red) or negative (blue) regression coefficient. The plots are dual-coded and simultaneously display the contrast estimate (x-axis) and t-values (y-axis). The hue indicates the size of the contrast estimate, and the opacity indicates the height of the t-value. The z-coordinates correspond to the standard MNI brain. The data are plotted using a procedure introduced by Allen et al. (2012) and implemented by Zandbelt, (2017).

First, there were no main effects of dopamine synthesis capacity on curiosity in the ventral striatum (**BRMS:** 95% CI [−.35, .32]), the caudate nucleus (**BRMS:** 95% CI [−.16, .51]) and the putamen (**BRMS:** 95% CI [−.35, .32]). Thus there was no evidence for a link between dopamine and overall curiosity ratings.

Furthermore, we found no interactions between outcome uncertainty and dopamine synthesis capacity on curiosity in the ventral striatum (**BRMS**: 95% CI [−.31, .31]), the caudate nucleus (**BRMS:** 95% CI [−.31, .31]) and the putamen (**BRMS:** 95% CI [−.30, .32]). There were also no interactions between outcome valence (gain/loss) and dopamine synthesis capacity on curiosity in the ventral striatum (**BRMS:** 95% CI [−.44, .037]) and the putamen (**BRMS:** 95% CI [−.44, .36]). However, we did find a significant interaction between outcome valence (gain/loss) and dopamine synthesis capacity on curiosity in the caudate nucleus (**BRMS:** 95% CI [−.51, −.056]), such that people with lower dopamine synthesis capacity were more curious about gains compared with losses than people with higher dopamine synthesis capacity. However, this effect was driven by four participants with unusually large differences between gain and loss trials (Figure 3A and Supplementary Figure 1A).

Also, there were no interactions between absolute expected value and dopamine synthesis capacity on curiosity in the ventral striatum (**BRMS:** 95% CI [−.18, .058]), the caudate nucleus (**BRMS:** 95% CI [−.084, .15]) and the putamen (**BRMS:** 95% CI [−.15, .088]). The same was true for the three-way interaction between absolute expected value, outcome valence (gain/loss) and dopamine synthesis capacity in the ventral striatum (**BRMS:** 95% CI [−.15, .022]), the caudate nucleus (**BRMS:** 95% CI [−.15, .020]) and the putamen (**BRMS:** 95% CI [−.17, .0069]).

## Discussion

We investigated whether curiosity ratings obtained from a non-instrumental lottery task vary with individual differences in dopamine synthesis capacity. We found no evidence that curiosity was related to participants’ dopamine synthesis capacity in the ventral striatum, the caudate or the putamen. Sensitivity to motives underlying curiosity, such as outcome uncertainty (Kobayashi et al., 2019; Romero Verdugo et al., 2020; van Lieshout et al., 2020a, 2018; van Lieshout, Traast, et al., 2019), outcome valence (Charpentier et al., 2018; Marvin & Shohamy, 2016; van Lieshout et al., 2020a; van Lieshout, Traast, et al., 2019) and expected value (Charpentier et al., 2018; Kobayashi et al., 2019; Romero Verdugo et al., 2020; van Lieshout, Traast, et al., 2019), were also not predicted by individuals’ dopamine synthesis capacity.

Previous work has linked dopamine synthesis capacity to ventral striatal coding of reward prediction errors (Deserno et al., 2015; Schlagenhauf et al., 2013; Boehme et al., 2015), reward-based reversal learning (Cools et al., 2009), cognitive control (Aarts et al., 2014) and cognitive effort (Hofmans, Papadopetraki, et al., 2020; Westbrook, van den Bosch, et al., 2020). Here we obtained no support for the hypothesis that these findings extend to a different form of cognitive motivation, namely non-instrumental curiosity. The absence of such a link may be surprising given that prior work in macaque monkeys using single-neuron recording (Bromberg-Martin & Hikosaka, 2009, 2011) and in humans using fMRI (Charpentier et al., 2018) demonstrated that the desire for knowledge in non-instrumental settings is implicated in the mesolimbic reward circuitry. Also, work with trivia questions has associated self-reported curiosity with brain activity in the caudate nucleus (Kang et al., 2009), the midbrain and the nucleus accumbens (Gruber et al., 2014). Similarly, receiving information has been associated with ventral striatum activity using trivia questions (Ligneul, Mermillod, & Morisseau, 2018) as well as in perceptual curiosity paradigms (Jepma, Verdonschot, van Steenbergen, Rombouts, & Nieuwenhuis, 2012). Here, we do not obtain support for the hypothesis raised by that prior work, by demonstrating that we have no evidence for a link between individual variability in non-instrumental curiosity and individual variation in dopamine synthesis capacity.

In a previous fMRI study using a similar lottery paradigm as the current study (van Lieshout et al., 2018), we found no evidence for activity in striatal areas as a function of induction or relief of curiosity. This might be due to this particular task not implicating the striatum or dopamine because of its passive nature, but it might also be explained by lower signal-to-noise ratio in deep brain structures due to the specific fMRI sequence used in that study (see van Lieshout et al., 2018). Similarly, one possible explanation for a lack of effects in the current study might be a too low signal-to-noise ratio of the [^18^F]DOPA radiotracer, which is a substrate for catechol-O-methyltransferase (COMT) in the periphery. As such, metabolites can cross the blood-brain-barrier and will distribute throughout the brain in a uniform fashion. This enhances background noise relative to the use of for example [^18^F]FMT, which is not a substrate for COMT (Becker et al., 2017), leading to a lower signal-to-noise ratio. It should be noted that this would mainly be a concern when the regions of interests are located in brain areas with low dopamine levels, but less so in the dopamine-rich striatum. Also, the risk of a too low signal-to-noise ratio was reduced by administering entacapone, which inhibits peripheral COMT metabolism, before PET scanning. [^18^F]DOPA and [^18^F]FMT also differ in their metabolic actions after decarboxylation by aromatic amino acid decarboxylase (AAAD), including higher affinity of [^18^F]DOPA metabolites compared with [^18^F]FMT metabolites for the vesicular monoamine transporter, leading to increased cell clearance of radiolabeled [^18^F]DOPA metabolites (Doudet et al., 1999). However, this would mostly be a concern for extended scanning times, as [^18^F]DOPA behaves as an irreversibly bound tracer in the first 90 minutes after tracer injection, during which their uptake rates are tightly correlated (Becker et al., 2017; Doudet et al., 1999).

The lack of a relationship between dopamine synthesis capacity and curiosity does not necessarily mean that dopamine transmission plays no role in curiosity. Dopamine levels in the brain are not a function of dopamine synthesis capacity alone, but also of other factors not measured in the current study (i.e. dopamine receptor availability, transporter density, dopamine release and genetic make-up). Thus, the current study does not refute hypothesized correlations between curiosity levels and other measures of dopamine function and stresses the need for further studies. For example, it might be the case that we find no correlation between curiosity and dopamine synthesis capacity per se, but that there would be dopaminergic drug effects as a function of dopamine. Here, dopamine synthesis capacity can be seen as a trait index of dopamine which itself is not correlated with curiosity, but does heighten the potential for phasic dopamine to have an effect (in agreement with adaptive gain theories looking at norepinephrine; Aston-Jones & Cohen, 2005).

Additionally, instead of using measuring dopamine synthesis capacity, using radioligands that bind to dopamine D2-receptors, such as raclopride or fallypride, may be interesting options for future research. These enable one to measure D2-receptor availability and, after a pharmacological challenge (e.g. methylphenidate), to measure dopamine release. The extra dopamine released after drug intake will compete with the radioligand for binding to D2 receptors. This reduction in PET signal as a result of the reduced receptor binding by the radioligand provides an index of dopamine release. Given the well-known role of the large ascending neuromodulators (i.e. dopamine and noradrenaline) in the various curiosity-relevant constructs highlighted here, such as uncertainty-based (meta-)learning (i.e. Nassar et al., 2012; Wang et al., 2018), reward motivation and cognitive effort (Hofmans, Papadopetraki, et al., 2020; Westbrook, van den Bosch, et al., 2020), human psychopharmacological interventions for studying the basis of both inter- and intra-individual variability in curiosity behavior might be promising.

Despite not finding a relationship between dopamine synthesis capacity and curiosity, the behavioral analyses provided a replication of our previous work. First of all, we demonstrate that curiosity is a function of outcome uncertainty, such that curiosity increased with increasing outcome uncertainty (Kobayashi et al., 2019; Romero Verdugo et al., 2020; van Lieshout et al., 2020a, 2018; van Lieshout, Traast, et al., 2019). Additionally, curiosity was greater for positive information (gains) compared with negative information (losses; Charpentier, Bromberg-Martin, & Sharot, 2018; Marvin & Shohamy, 2016; van Lieshout et al., 2020a; van Lieshout, Traast, et al., 2019). Again, we found no interaction between outcome uncertainty and outcome valence on curiosity, strengthening the claim that these factors operate largely independent from each other (see van Lieshout, Traast, et al., 2019).

To conclude, we find no evidence that individual variability in non-instrumental curiosity can be accounted for by individual variation in dopamine synthesis capacity. At the same time, the current study does not refute hypothesized correlations between curiosity levels and other measures of dopamine function and stresses the need for human psychopharmacological interventions for studying the basis of both inter- and intra-individual variability in curiosity.

## Acknowledgements

This work was supported by The Netherlands Organization for Scientific Research (NWO Vidi award 452-13-016 to FPdL and NWO Vici award 453-14-015 to RC), the James McDonnell Foundation (JSMF scholar award 220020328 to RC) and the EC Horizon 2020 Program (ERC starting grant 678286 awarded to FPdL). We thank Jessica I. Määttä for assistance with project administration and Felix Linsen for assistance during data collection

## Notes

### Competing Interest Statement

The authors have declared no competing interest.

