## Supplementary Material for "Does dopamine synthesis capacity predict individual variation in curiosity?"

**Supplement: Analyses using Repeated Measures ANOVAs**

In addition to the analyses reported in the main text, we performed similar analyses using repeated measures ANOVAs in SPSS (RRID:SCR_002865) and Bayesian repeated measures ANOVAs in JASP (RRID:SCR_015823). These analyses were performed to accommodate readers who are used to interpreting frequentist statistics instead of Bayesian credible intervals and to verify that our conclusions do not depend on the analytical framework employed.

To this end, we divided the values of outcome uncertainty into “low outcome uncertainty” and “high outcome uncertainty”, such that approximately 50% of the trials were indicated as being low outcome uncertainty (outcome uncertainty <= 200) and approximately 50% as high outcome uncertainty (outcome uncertainty > 200). Additionally, we divided the values of absolute expected value into “low absolute expected value” (expected value (absolute) < 50) and “high expected value” (expected value (absolute) > 50). Note that the trials with absolute expected value = 50 were omitted from the analyses. This was done because the values of absolute expected value were perfectly centered around expected value (absolute) = 50, precluding us to classify these trials as being either low or high absolute expected value

First, as main analyses we performed 2 (outcome valence: gain, loss) x 2 (outcome uncertainty: low, high) x 2 (expected value (absolute): low, high) repeated measures ANOVAs with “outcome valence (gain/loss)”, “outcome uncertainty (low/high)” and “absolute expected value (low/high)” as within-subject factors. “Dopamine synthesis capacity (Ki value)” was added as covariate (z-scored). The dependent variable was mean curiosity as indicated by the curiosity ratings. These main analyses were performed for the three striatal regions of interest separately (the caudate nucleus, the putamen and the ventral striatum/nucleus accumbens). We report the uncorrected p-values, but because all p-values should be Bonferroni-corrected for the number of striatal regions that are statistically tested (three), the p-values are deemed significant if they are smaller than .017 (= .05 / 3).

Next, if any of the interaction-effects with outcome valence (gain/loss) was statistically significant, we ran 2 (outcome uncertainty: low, high) x 2 (expected value (absolute): low, high) repeated measures ANOVAs with “outcome uncertainty (low/high)” and “absolute expected value (low/high)” as within-subject factors on the gain and loss trials separately. “Dopamine synthesis capacity (Ki value)” was added as covariate (z-scored). The dependent variable was mean curiosity as indicated by the curiosity ratings.

Additionally, as for the BRMS analyses (see main text), we performed a similar repeated measure ANOVA without adding dopamine synthesis capacity as covariate. These results are reported under “main task effects”.

**Results**

**Main task effects**

When analyzing the data with repeated measures ANOVAs, we also found a main effect of outcome uncertainty, such that participants were more curious about high compared with low outcome uncertainty (**RMA:** F(1,42) = 166.1, *p* = 3.47e-16, η_p_^2^ = .80, BF_incl_ = 6.03e+48). In addition, we found a main effect of outcome valence, such that participants were more curious about gains compared with losses (**RMA:** F(1,42) = 8.3, *p* = .006, η_p_^2^ = .17, BF_incl_ = 2.02e+5). However, there was no interaction between outcome uncertainty and outcome valence (**RMA:** F(1,42) = .082, p = .78, η_p_^2^ = .002, BF_incl_ = .17), indicating that these effects operate independent from each other.

There was also a main effect of absolute expected value (**RMA:** F(1,42) = 10.2, *p* = .003, η_p_^2^ = .20, BF_incl_ = 1.33), but this time we found no interaction between outcome valence and absolute expected value (**RMA:** F(1,42) = .69, *p* = .41, η_p_^2^ = .016, BF_incl_ = .18). However, when we did analyse the gain and loss trials separately, as we also did using the BRMS package reported in the main text, we found that participants were more curious about higher gains compared to lower gains (**RMA:** F(1,42) = 9.6, *p* = .004, η_p_^2^ = .19, BF_incl_ = 3.67), but that there was no effect of expected value on losses (**RMA:** F(1,42) = 3.86, *p* = .056, η_p_^2^ = .084, BF_incl_ = .59).

**Individual differences**

Next, we investigated whether the effects of outcome valence (gain / loss), outcome uncertainty and absolute expected value were a function of the dopamine synthesis capacity (Ki values) in the striatal regions of interest (ventral striatum, putamen and caudate nucleus).

First of all, there were no main effects of dopamine synthesis capacity on curiosity in the ventral striatum (**RMA:** F(1,41) = 3.86e-4, *p* = .98, η_p_^2^ = 9.42e-6), the caudate nucleus (**RMA:** F(1,41) = 1.1, *p* = .30, η_p_^2^ = .026) and the putamen (**RMA:** F(1,41) = .085, *p* = .77, η_p_^2^ = .002). This means that the overall curiosity ratings participants gave during the task were not a function of their dopamine synthesis capacity.

Furthermore, we found no interactions between outcome uncertainty and dopamine synthesis capacity on curiosity in the ventral striatum (**RMA:** F(1,41) = .089, *p* = .77, η_p_^2^ = .002), the caudate nucleus (**RMA:** F(1,41) = .68, *p* = .42, η_p_^2^ = .016) and the putamen (**RMA:** F(1,41) = .43, *p* = .52, η_p_^2^ = .010). In other words: the effect of outcome uncertainty on curiosity was not modulated by participants’ dopamine synthesis capacity. The same held for the interaction between outcome valence (gain / loss) and dopamine synthesis capacity on curiosity in the ventral striatum (**RMA:** F(1,41) = 2.0, *p* = .17, η_p_^2^ = .046) and the putamen (**RMA:** F(1,41) = 3.3, *p* = .077, η_p_^2^ = .074). It should be noted here that we found a significant interaction effect between dopamine synthesis capacity and outcome valence on curiosity in the caudate nucleus (**RMA:** F(1,41) = 6.6, *p* = .014, η_p_^2^ = .14). However, this effect was driven by four participants with unusually large differences between gain and loss trials (Figure 3A and Supplementary Figure 1A).

Also, there were no interactions between absolute expected value and dopamine synthesis capacity on curiosity in the ventral striatum (**RMA:** F(1,41) = 1.8, *p* = .18, η_p_^2^ = .043), the caudate nucleus (**RMA:** F(1,41) = .24, *p* = .63, η_p_^2^ = .006) and the putamen (**RMA:** F(1,41) = 1.3, *p* = .26, η_p_^2^ = .031). The same was true for the three-way interaction between absolute expected value, outcome valence (gain/loss) and dopamine synthesis capacity in the ventral striatum (**RMA:** F(1,41) = 1.0, *p* = .32, η_p_^2^ = .024), the caudate nucleus (**RMA:** F(1,41) = 1.7, *p* = .20, η_p_^2^ = .040) and the putamen (**RMA:** F(1,41) = 3.1, *p* = .088, η_p_^2^ = .069).

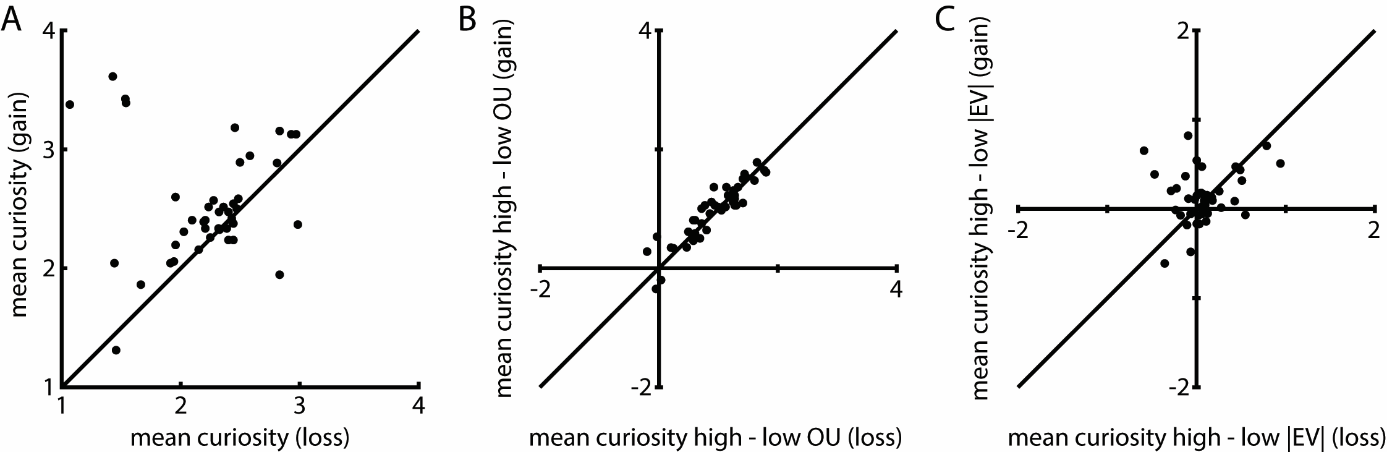

**Supplementary Figure 1. Behavioral results of the experiment – individual differences.­­­**

**A.** Panel A depicts the individual variability in the extent to which participants were more curious about gains compared with losses. The x-axis depicts mean curiosity for the loss trials and the y-axis the mean curiosity for the gain trials. Every dot represents one participant. The majority of the dots are above the diagonal, indicating that most participants were more curious about gains compared with losses.

**B.** Panel B depicts the individual variability in the extent to which participants were driven by outcome uncertainty for gain and loss trials. The x-axis depicts mean curiosity for high minus low outcome uncertainty in loss trials, with positive and negative values indicating a positive or negative relationship between outcome uncertainty and curiosity respectively. The y-axis depicts mean curiosity for high minus low outcome uncertainty in gain trials, with positive and negative values indicating a positive or negative relationship between outcome uncertainty and curiosity respectively. Every dot depicts one participant. The majority of participants have a positive relationship between outcome uncertainty and curiosity for both gain and loss trials.

**C.** Panel C depicts the individual variability in the extent to which participants were driven by absolute expected value for gain and loss trials. The x-axis depicts mean curiosity for high minus low absolute expected value in loss trials, which positive and negative values indicating a positive or negative relationship between absolute expected value and curiosity respectively. The y-axis depicts mean curiosity for high minus low absolute expected value in gain trials, which positive and negative values indicating a positive or negative relationship between absolute expected value and curiosity respectively. Every dot depicts one participant. The majority of participants have positive values on the y-axis, indicating that they are more curious about high compared with low gains. However, there is more variability in positive and negative values on the x-axis. As such, there was no effect of absolute expected value on curiosity for losses.
